# siRNADesign: A Graph Neural Network for siRNA Efficacy Prediction via Deep RNA Sequence Analysis

**DOI:** 10.1101/2024.04.28.591509

**Authors:** Rongzhuo Long, Ziyu Guo, Da Han, Xudong Yuan, Guangyong Chen, Pheng-Ann Heng, Liang Zhang

## Abstract

The clinical adoption of small interfering RNAs (siRNAs) has prompted the development of various computational strategies for siRNA design, from traditional data analysis to advanced machine learning techniques. However, previous studies have inadequately considered the full complexity of the siRNA silencing mechanism, neglecting critical elements such as siRNA positioning on mRNA, RNA base-pairing probabilities, and RNA-AGO2 interactions, thereby limiting the insight and accuracy of existing models. Here, we introduce **siRNADesign**, a Graph Neural Network (GNN) framework that leverages both non-empirical and empirical rules-based features of siRNA and mRNA to effectively capture the complex dynamics of gene silencing. In multiple internal datasets, siRNADesign achieves *state-of-the-art* performance. Significantly, siRNADesign also outperforms existing methodologies in *in vitro* wet lab experiments and an externally validated dataset. Additionally, we develop a new data-splitting methodology that addresses the data leakage issue, a frequently overlooked issue in previous studies, ensuring the robustness and stability of our model under various experimental settings. Through rigorous testing, siRNADesign has demonstrated remarkable predictive accuracy and robustness, making significant contributions to the field of gene silencing. Furthermore, our approach in redefining data-splitting standards aims to set new benchmarks for future research in the domain of predictive biological modeling for siRNA.

## Introduction

RNA interference (RNAi) serves as a critical regulatory mechanism in cells, utilizing short double-stranded RNA (dsRNA) molecules to guide the homology-dependent regulation of gene expression [1]. This process involves small interfering RNAs (siRNAs), which are RNAi-based regulators consisting of 19-23 nucleotides. These siRNAs are processed from longer dsRNA precursors by Dicer, an RNase III-like enzyme, and then incorporated into the RNA-induced silencing complex (RISC). Within RISC, siRNA strands are separated, and the strand with a more stable 5’ end guides the complex to bind homologously to the target mRNA. The catalytic component of RISC, a member of the argonaute family (AGO2), then cleaves the target mRNA, leading to its degradation and the silencing of the gene [2].

Thousands of potential siRNAs can target the same gene, thus, identifying the most effective siRNA from these candidates is currently a focus of study. Researchers have developed various algorithms to predict siRNA activity, categorized into **two** generations. The **first**-generation tools were based on empirical rules from validated small siRNA datasets, considering several features, such as GC content, base preference at specific positions, thermodynamic stability, internal structure, and the secondary structure of the target mRNA [3]. However, first-generation methods were shown to have low accuracy, with experiments indicating that about 65% of the predicted active siRNAs failed to achieve 90% knock-down efficacies [4]. Furthermore, approximately 20% of the siRNAs were found to be inactivated [4]. This issue could be attributed to the limited size of early datasets, which likely did not encompass many critical features [5]. With the expansion of siRNA data and the development of artificial intelligence, the **second**-generation algorithms have improved predictions of siRNA knockout efficiency by utilizing advanced data mining techniques. Huesken *et al*. [6] constructed a dataset containing 2431 siRNAs using a high-throughput analysis technique, and developed a tool named Biopredsi to predict siRNA efficiency, based on artificial neural networks [6]. Subsequently, with the contribution of Huesken’s dataset, numerous machine learning-based siRNA activity prediction tools emerged. DSIR used a simple linear LASSO model combining basic features of siRNA sequences for siRNA efficacy prediction with a good performance [7]. Additionally, i-Score algorithm [8] predicted active siRNAs by applying a linear regression model, only comprised of nucleotide (nt) preferences at each position without other parameters. However, **two main issues** impair the performance of these early second-generation algorithms. ***Firstly***, the performance of these algorithms exhibits instability, potentially stemming from the heterogeneity of siRNA datasets, which are assessed by various groups across different protocols, and largely depends on feature selection [9]. ***Secondly***, the numerous interactive factors and nonlinear characteristics of biological characteristics in biological processes significantly challenge the construction and optimization of traditional models [10].

To address the first issue, some advanced artificial intelligence models have been employed in the development of second-generation algorithms. For instance, Han *et al*. [11] introduced deep learning scenarios, i.e., Convolutional Neural Network (CNN), into this field to achieve a more powerful siRNA efficacy predictor. Specifically, they utilized convolution and linear layers to extract hidden features from sequence context and thermodynamic properties, followed by a deep architecture for efficacy prediction. This CNN method was proven to be more stable and efficient than previous works [8, 12] for the prediction of siRNA silencing efficacy. Although, CNN model showed advantages in addressing the first issue, it struggled to consider the numerous interactive factors and nonlinear characteristics of biological processes. Graph Neural Network (GNN) are able to analyze the hidden relationships and information within the data as CNN models, offering a straightforward and intuitive representation of heterogeneous and complex biological processes [13]. In contrast, GNN provides superior handling of the second issue concerning internal interaction networks within the data. As an effective solution for multi-modal information fusion and exploration, GNN excels in analyzing multi-modal data represented in graphical structures, including the language of sequences and biological networks [14]. In biological molecular networks, siRNA and mRNA constitute a complex interaction network, where each mRNA molecule may interact with numerous different siRNA molecules. Thus, GNN offers a potent solution for constructing a siRNA-mRNA interaction network to predict their interaction ability based on the features of siRNA and mRNA.

To enhance the understanding of the siRNA silencing process using GNN model, incorporating specific features crucial features that impact siRNA silencing efficiency is imperative. Different type of nodes benefits from the integration of relevant features to better model the complexities of RNA interference. Previous research reported that the sequence of siRNA and mRNA influenced the efficacy of RNA interference [15]. Additionally, the prevailing view showed that the thermodynamic stability profile of the siRNA duplex strongly influenced the efficacy of siRNA by reflecting a guide strand selection mechanism [16]. Therefore, recently, Massimo *et al*. [17] introduced the first GNN-based model named GNN4siRNA for siRNA inhibition prediction, featuring three node types: siRNA and mRNA nodes with k-mer counts, and siRNA-mRNA interaction nodes with features from thermodynamic parameters. The features of interaction edges were attributed to interaction nodes, while the weights of the original interaction edges served as labels for these nodes. Utilizing the proposed graph topology enabled the prediction of siRNA efficiency through node regression tasks.

However, k-mer counts used in GNN4siRNA, focusing on local sequence patterns with multiple nucleotides, potentially overlooked or diminished specific nucleotides positions. Additionally, inappropriate k-values may lead to omission of crucial information or introduction of noise. Moreover, there are several key factors previously discussed that critically influence the efficacy of siRNA-mediated gene silencing ignored by GNN4siRNA. For example, the positional context of siRNA along the mRNA strand was crucial for its gene silencing efficacy [18]. What’s more, previous studies indicated that the base-pairing secondary structure was more crucial for RNA’s proper function than the primary sequence itself in many cases [19]. Importantly, many empirical rules as the key point of first-generation tools, including nucleotide frequencies [20], G/C percentages of siRNA [21], and the impact of nucleotides at each position within siRNA [22], were excluded by GNN4siRNA. Therefore, GNN4siRNA’s mining of sequence characteristics was not in-depth or broad enough, which did not account for several key factors previously discussed that critically influence the efficacy of siRNA-mediated gene silencing, undermining the performance of model.

To this end, we propose **siRNADesign**, a GNN framework that thoroughly explores the sequence features of siRNA and mRNA with a specific topological structure, enhancing siRNA efficacy prediction performance. Our siRNADesign firstly extracts two distinct-type features of RNA, i.e., non-empirical features and empirical-rule-based ones, and integrates them into GNN training. Specifically, the non-empirical features include one-hot sequence encoding, position encoding, base-pairing probabilities, and RNA-protein interaction probabilities. The empirical-rule-based features include the thermodynamic stability profile, nucleotide frequencies, the G/C percentages, and the rule codes in each position of siRNA. Compared with other existing methods, the experiment results on all datasets demonstrate that our model achieves *state-of-the-art* performance.

Upon that, we notice a data leakage issue within the cross-validation setting, i.e., a mRNA from the test set might have been included in the training phase, thereby compromising the integrity of the evaluation. Therefore, we further propose **a new splitting method for evaluation** that randomly divides the dataset into three subsets for training, evaluation, and testing. To avoid data leakage or deviation from randomness, we respectively treat siRNA and mRNA as the basis of division and repeat multiple times with different seeds. This evaluation setting enables a more rational and effective evaluation of models’ predictive performance and robustness. We conduct extensive experiments on internal datasets and validation on external datasets, including *in vitro* wet lab experiments and an externally collected dataset. Our siRNADesign shows superior performance on all datasets under different splitting settings.

The main contributions of our paper are summarized as follows:

- We propose **siRNADesign**, a GNN framework that thoroughly explores the sequence features of siRNA and mRNA, and boosts siRNA efficacy prediction performance.
- To tackle the data leakage issue in the previous evaluation settings, we propose **a new splitting method for evaluation** that achieves a more rational and effective evaluation of models’ predictive performance and robustness, serving as a new benchmark for future works.
- We conduct experiments on the common-used dataset [17] under our proposed evaluation settings, where siRNADesign achieves *state-of-the-art* performance.
- We conduct *in vitro* **wet lab** studies and also collect a dataset for external validation, and our model shows superior performance on both datasets compared with others across all metrics.

## Materials and Methods

In this section, we elaborate on the proposed data splitting manner and model details of siRNADesign.

### Data sources

#### Public data

In this work, we collected the siRNA datasets with 2816 siRNA-mRNA pairs and related efficacy values from Massimo *et al*. [17], which came from the original studies of Huesken [6], Harborth [23], Ui-Tei [24], Vickers [25], and Khovorova [26]. We termed this dataset as ‘Dataset HUVK’. Additionally, we utilized the Simone dataset [27], which comprised 322 siRNAs, to further validate our models.

#### In-house data

What’s more, we conducted *in vitro* wet lab studies, and the collecting procedures were followed to test the silencing efficiency of the designed siRNA on the target SERPINC1. In detail, on day 0, 293A cells (ATCC, Manassa, VA, US) were cultured in a 96-well cell culture plate with 100 *µ*L culture medium. The initial seeding was 12,000 cells per well. On day 2, the medium was replaced with 150 *µ*L. 0.4 *µ*L of Lipofectamine 2000 (Thermo Fisher Scientific, Waltham, MA, US) was used as a plasmid transfection agent for the cells. Four different concentrations (10 nM initial, 10-fold dilution) of designed siRNA (in-house synthesized by ACON Pharmaceuticals, Cranbury, NJ, US), the negative control wells (transfection with luciferase plasmid only) and the background control wells (wells containing cells) were used. On day 2, after 24-hour transfection, double luciferase detection was performed, and silencing efficiency was calculated accordingly.

We conducted multiple trails for each siRNA and obtained silencing efficiencies for 298 siRNAs. Those demonstrating negative knockdown efficiency were removed from further consideration. From the remaining siRNAs, we eliminated those that exhibited significant variability across trials due to experimental errors. By computing the average knockdown efficacy across various trials within the same experimental setup, we refined our dataset to include 102 siRNAs with consistent results, forming our final validation set.

In Table 1, we summarized the characteristics of the three datasets utilized in our experiments, including the number of mRNAs, the number of siRNAs, and the count of siRNA-mRNA pairs.

**Table 1.** Dataset Information. We summarize the information of the three datasets utilized in our experiments, including mRNA number, siRNA number, and siRNA-mRNA pair number.

| Dataset Name | siRNA Num. | mRNA Num. | Pairs Num. |
| --- | --- | --- | --- |
| Dataset_HUVK [17] | 2816 | 44 | 2816 |
| Simone [27] | 322 | 6 | 322 |
| In-house(ACON) | 102 | 1 | 102 |

### Dataset Splitting

Prior works [8, 11, 12, 17] evaluated the models via 10-fold cross-validation. Specifically, each dataset was divided into ten equal parts, with nine parts used for training and the remaining one part for testing. This cycle was executed ten times, allowing each segment to serve as the test set once. However, we observed a data leakage issue within the cross-validation setting, i.e., a siRNA from the test set might have been included in the training phase, thereby compromising the integrity of the evaluation. What’s more, existing methods [8, 11, 12, 17] employed the same subset for both validation and testing, which was not reasonable and is impractical for real-world applications. Consequently, we introduced a **new** evaluation setting. Here, we randomly divided the dataset into a training set, an evaluation set, and a test set with a ratio of 70:15:15. To avoid data leakage or deviation from randomness, we respectively treated siRNA and mRNA as the basis of division and repeat the splitting process 10 times with different random seeds. This approach yielded 20 distinct splits, offering a more robust and realistic evaluation. Note that we denoted the split datasets divided based on siRNA and mRNA as ‘siRNA-split data’ and ‘mRNA-split data’ respectively.

### Preliminaries: GNNs and GCNs

In this section, we delve into the design and utility of classical Graph Neural Networks (GNNs) and Graph Convolutional Networks (GCNs), which form the foundational elements of our model tailored for processing graph-structured data.

Unlike conventional machine learning models that operate in Euclidean spaces, GNNs are uniquely adept at managing data that naturally forms networks or graphs. This is particularly relevant in the biological research sphere, where complex systems like protein interactions and metabolic pathways are often conceptualized as networks. GNNs mathematically represent graphs as *G* = (*V, E*), with *V* denoting vertices or nodes, and *E* representing edges where each edge *e*_*ij*_ ∈ *E* connects nodes *v*_*i*_ and *v*_*j*_. The structure of a graph is further detailed by its adjacency matrix *A* ∈ ℝ^*n×n*^, while node and edge attributes are depicted through feature matrices *X*_*v*_ ∈ ℝ^*n×d*^ and *X*_*e*_ ∈ ℝ^*e×c*^, accommodating heterogeneity among nodes with distinct features.

Building upon the GNN framework, GCNs integrate the convolutional concept traditionally used in image processing to enhance the analysis of graph-structured data. By aggregating feature information from a node’s local neighborhood through the adjacency matrix *A* and the node and edge feature matrices *X*_*v*_ and *X*_*e*_, GCNs capture the topological structure of the graph. This aggregation enables GCNs to learn node representations that incorporate both their features and the influences of their adjacent nodes. The capability of GCNs to encode both node and structural information into dense vector representations proves highly effective for tasks such as node classification, link prediction, and graph classification, where spatial relationships among data points are critical.

### Model Structure

As shown in Fig 1, given the input siRNA and mRNA sequences, our model first extracts sequence characteristics deeply, and then assigns the features to the corresponding GNN nodes for feature fusion and efficacy regression. In the following paragraphs, we introduce the RNA sequence features and model structure in detail.

**Fig. 1.**
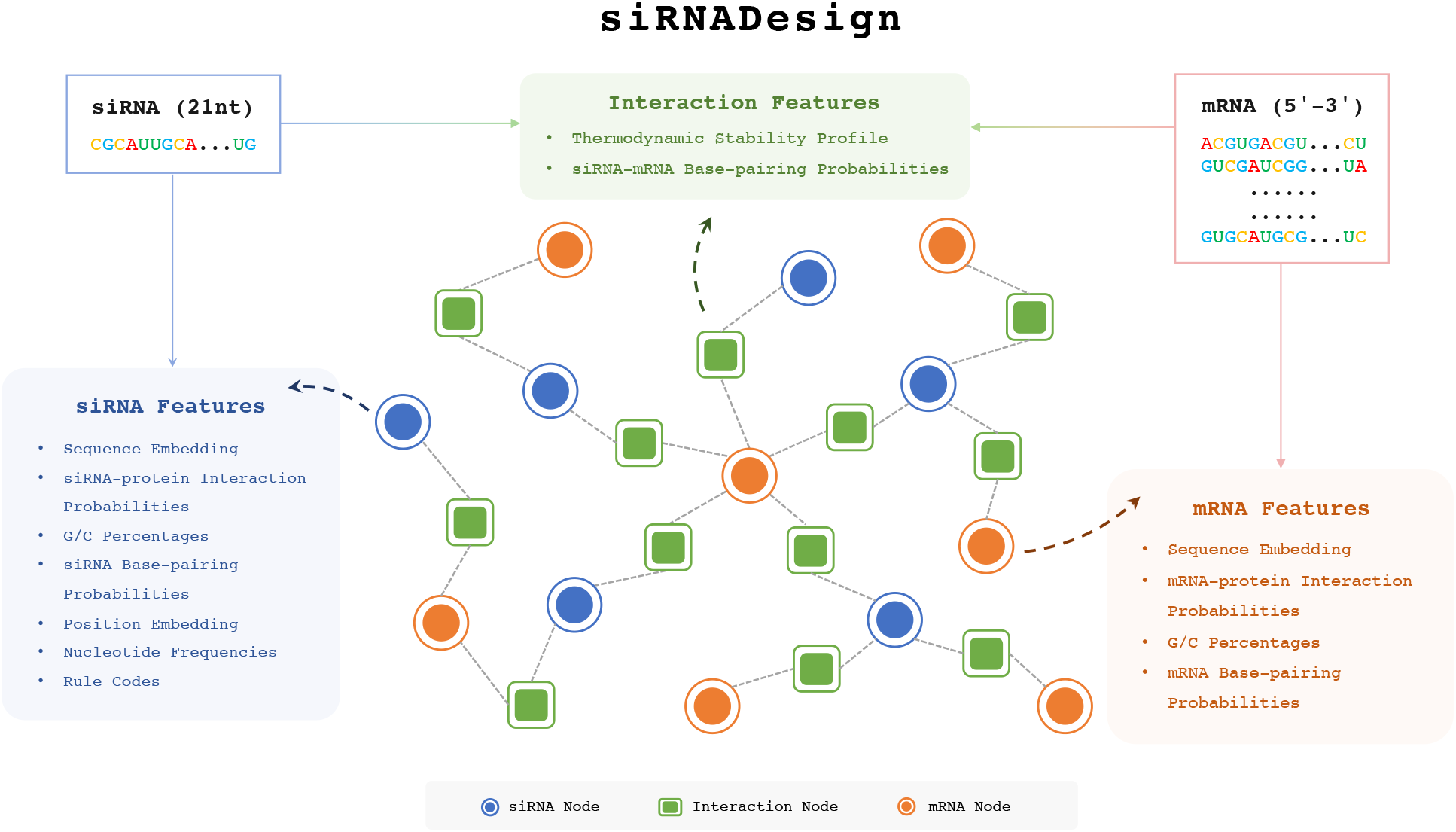
Overall Works of siRNA-DESIGN. Our model contains two modules, i.e., a **feature extraction module** for deep sequence characteristics exploration, and a **GNN prediction module** for feature fusion and efficacy regression.

## GNN Node Feature Assignment

To fully grasp the information embedded within mRNA and siRNA sequences, we encompassed the initial extraction of **two** types of features from the input RNA sequences and then assigned the features to the nodes of our GNN model. **The first type comprised non-empirical rule features**, which included sequence embeddings, positional embeddings, siRNA-mRNA base-pairing probabilities, and RNA-protein interaction probabilities. **The second type was constituted by empirical rule features**, which were thermodynamic stability profile, nucleotide frequencies, G/C percentages, and siRNA rule codes. To further explore the intrinsic properties of mRNA/siRNA and their interaction, we employed advanced methodologies [28, 29, 30, 31] to extract the aforementioned features. These methodologies collectively formed the feature extraction phase of our method.

Our model contained three types of nodes, i.e., the mRNA nodes, siRNA nodes, and the siRNA-mRNA interaction nodes, which were introduced in detail in Section 2.4. In Table 2, we showed the features assigned to each kind of node respectively. Below, we presented the non-empirical rule features and empirical rule features in detail.

**Table 2.**
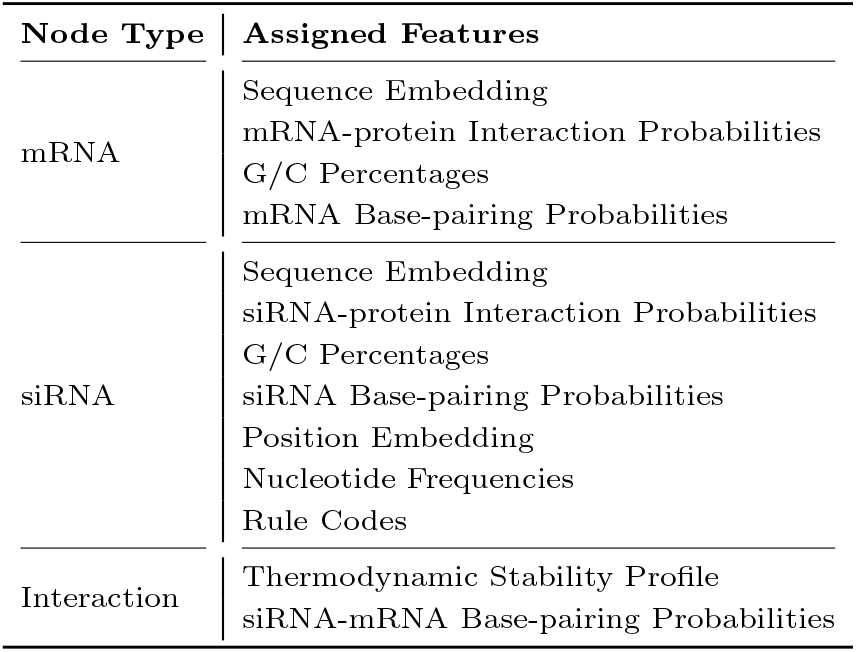
Assigned Features for Different Nodes. We show the features assigned to each kind of node. ‘mRNA’, ‘siRNA’, and ‘Interaction’ represent the mRNA node, siRNA node, and the siRNA-mRNA interaction node respectively.

| Node Type | Assigned Features |
| --- | --- |
| mRNA | Sequence Embedding<br>mRNA-protein Interaction Probabilities<br>G/C Percentages<br>mRNA Base-pairing Probabilities |
| siRNA | Sequence Embedding<br>siRNA-protein Interaction Probabilities<br>G/C Percentages<br>siRNA Base-pairing Probabilities<br>Position Embedding<br>Nucleotide Frequencies<br>Rule Codes |
| Interaction | Thermodynamic Stability Profile<br>siRNA-mRNA Base-pairing Probabilities |

### For the non-empirical rule features

#### Sequence Embedding

We employed one-hot encoding to represent biological sequences. Each nucleotide of the siRNA and mRNA sequences, corresponding to the four DNA bases A, C, G, T, was encoded by a unique four-dimensional binary vector: A = *<*1, 0, 0, 0*>*, C = *<*0, 1, 0, 0*>*, G = *<*0, 0, 1, 0*>*, T = *<*0, 0, 0, 1*>*. For ambiguous nucleotides, we denoted them as ‘N’ with a distinct vector *<*0, 0, 0, 0*>*.

#### Positional Embedding of siRNA

The positional context of siRNA along the mRNA strand was crucial for its gene silencing efficacy [18]. Each positional information of siRNA along the related mRNA is encoded into a higher-dimensional space of dimension *D*, then a positional encoding matrix **P** of a dimension *n × D* is calculated in an element-wise manner. **P**_*ij*_ is defined as below:

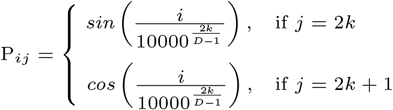

Where *i* denotes the position of siRNA on the mRNA strand and *j* is the channel dimension.

#### siRNA-mRNA Base-pairing Probabilities

The previous research reported that the efficacy of RNA interference(RNAi) approaches was primarily determined by the sequence of siRNA or the structure of mRNA [15]. Both siRNA and mRNA strands were capable of folding into secondary structures independently, a process largely by nucleotide base pairing including canonical (A-U, C-G), non-Watson-Crick (G-U), and non-canonical interactions, establishes its secondary structure [32]. Previous studies indicated that the base-pairing structure was more crucial for RNA’s proper function than the primary sequence itself in many cases [19]. Additionally, siRNA and mRNA could form complexes through intermolecular base pairing. Consequently, we used the RNAfold tool to obtain the base-pairing probability matrix for siRNA and mRNA, and RNAcofold for the siRNA-mRNA interaction base-pairing probability matrix. Both tools were part of the ViennaRNA package [28]. Due to the prevalence of zero probabilities in each matrix, truncated singular value decomposition (SVD) was employed to reduce dimensions and serve as a feature of the model.

#### RNA-protein Interaction Probabilities

In the process of RISC assembly, the guide strand binds to the Argonaute protein AGO2 and directs RISC to its complementary target RNA, which is subsequently cleaved and degraded by the RNase activity within AGO2 [33]. RNA-protein interactions are significant in the RNAi process, a factor previously overlooked in earlier studies for siRNA efficacy prediction. RPISeq [29] was a machine learning-based tool designed to predict RNA-protein interactions. It required sequences of an RNA and a protein as input, and provided interaction probability classifiers through two methods: RPISeq-SVM(Support Vector Machine) and RPISeq-RF (Random Forest). We utilized this tool to predict the interaction between siRNA-AGO2 and mRNA-AGO2, obtaining probabilities using RPISeq-RF as a feature of our model.

### For the empirical rule features

#### Thermodynamic Stability Profile

The prevailing view showed that the thermodynamic stability profile of the siRNA duplex strongly influenced the efficacy of siRNA by reflecting a guide strand selection mechanism [16]. The thermodynamic stability profile of the siRNA antisense strand included the calculation of Watson–Crick pair free energy (Δ*G*) between every two adjacent nucleotides, the total energy of the siRNA and the difference of energy at the 5’ and 3’ end of siRNA, all these computations followed the work [30].

#### Nucleotide Frequencies

Certain short motifs influenced the function of siRNA. For instance, sequences containing AAAA or UUUU were likely to be terminated by RNA polymerase III transcriptions, and CCCC or GGGG motifs may impact RNAi function by forming a nucleotide quartet [20]. Nucleotide frequency represented the count of each nucleotide within a siRNA sequence, and it was widely adopted in existing research [9, 34, 35]. Given the constant length of siRNAs, we calculated the frequencies of 1-mer (A, U, C, G), 2-mer (e.g., AU, CG, etc), 3-mer (e.g., AUC, UCG, etc), 4-mer (e.g., AUCG, UCGA, etc) and 5-mer (e.g., AUCGA, UCGUA, etc) segments with 4, 16, 64, 256, 1024 possible motifs, respectively.

#### G/C Percentages

GC content was a critical factor in the efficacy of siRNAs. Low GC content led to weak and non-specific binding, conversely, high GC content could hinder the unwinding of the siRNA duplex by RSIC complex and helicase [21]. Moreover, the past study indicated that lower GC content in both the global and local flanking regions of siRNA binding sites led to siRNA inhibition. Thus, we computed the GC content of siRNA and mRNA sequences [16].

#### Rule Codes of siRNA

Previous studies have shown that certain nucleotides at specific positions could either enhance or impair the function of siRNA. Initially, the impact of nucleotides at each position within siRNA was cataloged in the paper [22] and subsequently simplified by He *et al*. [31]. This effort was made to clarify the situation, as different rules explain preferences for specific positions in siRNA. In the simplified rules proposed by He *et al*., an encoding of 1 indicated a nucleotide’s preference for enhancing siRNA efficiency, whereas an encoding of -1 denoted a preference for reducing efficiency. If no rule specifies such a preference, the encoding defaulted to 0. Here, we represented these rules using a unique three-dimensional binary vector: 1 = *<*0, 0, 1*>*, 0 = *<*0, 1, 0*>*, -1 = *<*1, 0, 0*>*.

## GNN Model of siRNADesign

We modeled the graph of siRNADesign as an undirected heterogeneous graph with three distinct node types, i.e., siRNA, mRNA, and their interactions. We approached efficacy prediction as a node regression task, where outputs from the feature extraction module were aptly assigned to corresponding nodes. The strategy for assigning extracted features was detailed in Table 2. As shown in Figure 1, *M*_*j*_, *j* ∈ [1, *m*] and *S*_*i*_, *i* ∈ [1, *s*] were involved within the constructed graph, and *m, s* were the total number of mRNA and siRNA respectively.

Also, node 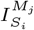 appeared in the graph, indicating known interaction features to the GNN module. Following [17], we adopted the Heterogeneous GraphSAGE platform [36], which utilized a dual-layer structure to aggregate each node’s feature with features of its neighbors. For any given node *v*, we utilized *N* (*v*) to represent its neighborhood, facilitating the aggregation process. The feature vector of node *v* at layer *l* was denoted by 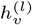, while 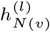 represented the aggregated feature vector from its neighborhood, the update rule for node *v* is formalized as:

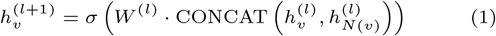

where *W* ^(*l*)^ was a layer-specific weight matrix, *σ* was a non-linear activation function, and CONCAT denotes the concatenation of the node’s current features with the aggregated neighborhood features.

### Hyperparameter Tuning

To avoid overfitting, we employed Optuna [37], a Bayesian optimization library, for efficient hyperparameter tuning and benchmarking. Optuna was an advanced framework specifically designed to facilitate the automatic exploration of hyperparameter spaces, aiming to identify the most effective combinations of hyperparameters through a systematic approach. In our study, we performed 100 trials of Bayesian optimization, adopting the Pearson Correlation Coefficient (PCC) as the primary metric to be maximized.

### Model Evaluation

To evaluate the efficacy of our model, we employed three key metrics: the Pearson Correlation Coefficient (PCC), the Spearman Correlation Coefficient (SPCC), and the Area Under the receiver operating characteristic Curve (AUC). Both PCC and SPCC served to measure the correlation between the actual and predicted efficacies, with PCC applied to continuous data and SPCC to ordinal rankings. The formulas for these coefficients were as follows:

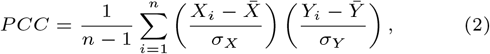

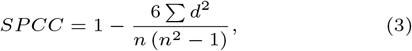

where *X* and *Y* denoted the predicted values and observed labels, and *n* was their common size. *d* represented the difference between the *X* rank and *Y* rank for each pair of data. In addition to these correlation coefficients, the AUC metric was employed to measure the overall predictive performance of the model. AUC values ranged from 0 to 1, where higher values indicated better performance.

## Results

### Training Settings and Model Settings

We adopted Oputna [37] to conduct hyperparameter tuning. We encompassed a series of trials focusing on various parameters, including the sizes of HinSAGE layers, hop neighbor samples, batch sizes, dropout rates, dimensions of positional embeddings, and the dimensions of reduced matrices for both RNA base-pairing probabilities and siRNA-mRNA base-pairing probabilities. The final settings and configurations of siRNADesign on siRNA-split data are comprehensively outlined in Table 3.

**Table 3.** Trainging Hyperparameters and Model Hyperparameters of siRNADesign <u>on siRNA-split data</u>. We list the best combination of the training and model hyperparameters among all trails. ‘Dim.’ denotes feature dimension.

| Name | Values |
| --- | --- |
| **Training Hyperparameter** |  |
| Batch Size | 64 |
| Learning Rate | 1e-3 |
| Loss Function | MSE |
| Epoch Number | 26 |
| **Model Hyperparameter** |  |
| HinSAGE Layer Size | [64, 32] |
| Hop Neighbor Sample | [12, 6] |
| Dropout Rate | 0.1 |
| Dim. of Positional Embeddings | 6 |
| Dim. of mRNA Reduced Matrices | 100 |
| Dim. of siRNA Reduced Matrices | 6 |
| Dim. of Pair Reduced Matrices | 50 |

### Performance

To verify the effectiveness of our proposed method, we compared siRNADesign with five previous works, i.e., GNN4siRNA [17], DSIR [12], s-Biopredsi [8], i-Score [8] and CNN model [11]. We obtained the predicted results of DSIR, s-Biopredsi, and i-Score from the i-Score website. In addition, we reproduced the results of the CNN model [11] according to the introduction and details in its paper.

#### Performance on Dataset HUVK

We conducted experiments and reported the performances on siRNA-split data in Figure 2. Experiment results under mRNA-splitting settings were presented in Supplementary Materials. As presented in Section 2.2, to avoid deviation from randomness, we conducted experiments under 10 distinct splits divided under different random seeds and report the average metrics and variance values. As shown in Figure 2, our **siRNADesign outperformed the other five methods with small variation**, reaching an average PCC of 0.770, SPCC of 0.771, MSE of 0.020, and AUC of 0.874. Meanwhile, GNN4siRNA achieved an average PCC of 0.679, SPCC of 0.673, MSE of 0.026, and AUC of 0.821. Results of DSIR, s-Biopredsi and i-Score were worse than GNN4siRNA, while the CNN model reached the worst with an average PCC, SPCC, MES, and AUC of 0.393, 0.399, 0.050, and 0.703, respectively.

**Fig. 2.**
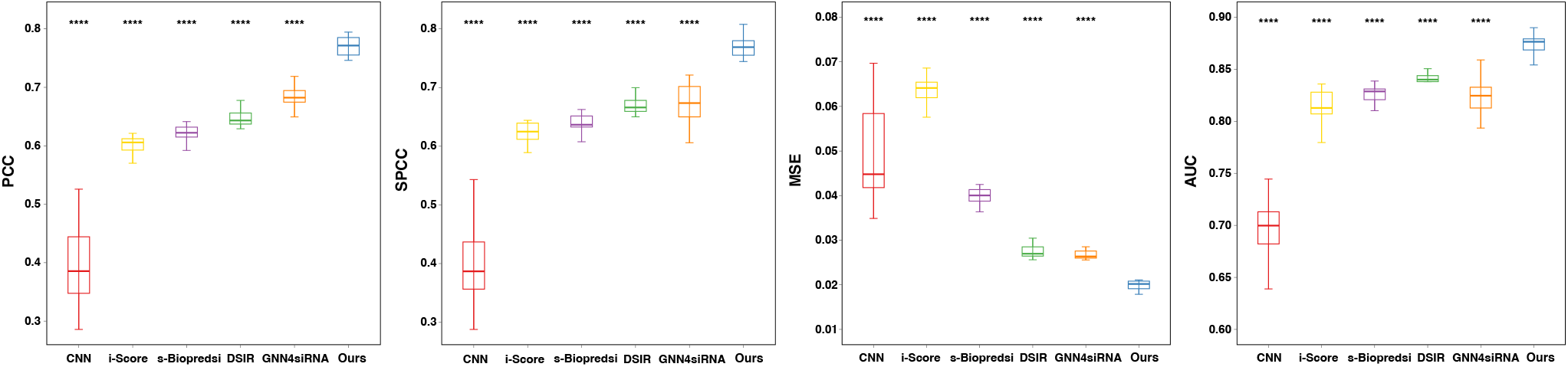
Performance of siRNADesign on siRNA-split Data (Dataset HUVK). Our method achieves superior performance and remarkable stability across multiple metrics over other models. PCC: Pearson correlation coefficient; SPCC: Spearman correlation coefficient; AUC: Area Under Curve; MSE: Mean squared error. P values are calculated using paired t-test to compare the siRNA_design with the metric of other models. ∗ ∗ ∗ ∗ *P* ≤ 0.0001.

#### Performance on Simone Dataset

Furthermore, we evaluated the effectiveness and robustness of these models on an external dataset, Simone [27], which contained 322 siRNAs. We first trained all models on the training set of siRNA-split data and utilized the pre-trained models to conduct inference on Simone. In Figure 3, we showed the evaluation results, where our siRNADesign showed the best performance across all metrics, achieving a PCC of 0.464, SPCC of 0.321 and AUC of 0.702. However, the CNN model’s predictions exhibited significantly lower performance, with a PPC of -0.017, SPCC of -0.054, and AUC of 0.4999, which were therefore omitted from the figure. It further demonstrated the superior robustness of our method even for external datasets compared with other methods.

**Fig. 3.**
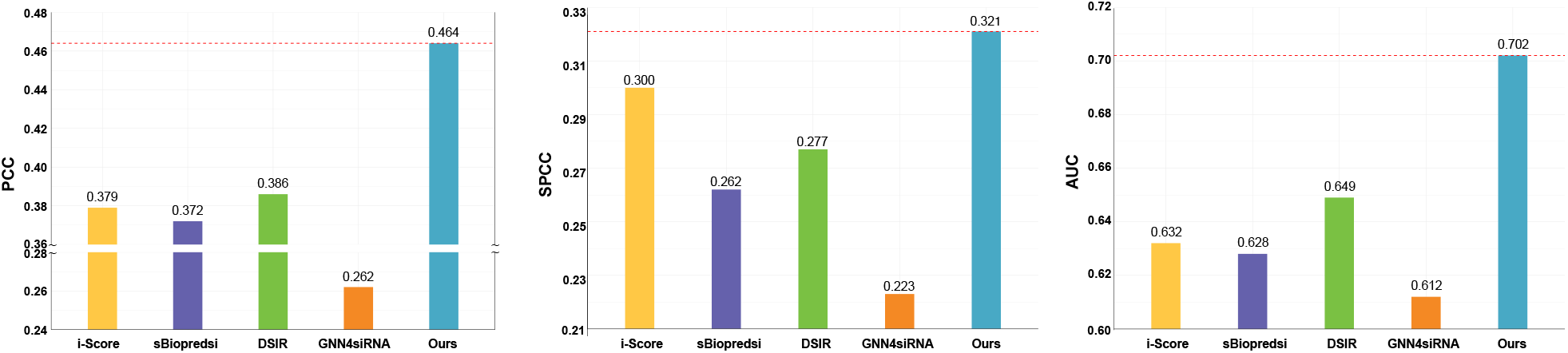
Performance of siRNADesign on an External Dataset, i.e., Simone [27]. Notably, our siRNADesign significantly surpasses other methods by large performance gains, demonstrating the strong efficacy prediction capacity of our model.

#### Result Analysis for Dataset HUVK and Simone

The unsatisfactory performance of DSIR, s-Biopredsi, and i-Score could be attributed to their reliance on traditional machine learning methods, which utilized feature engineering and depend on empirical rules. Furthermore, important features like the location of the siRNA within the mRNA and the information about interactions between the siRNA and mRNA were absent from the GNN4siRNA model. Its capacity to identify underlying patterns among the features was hampered by this omission. Meanwhile, the inferiority of the CNN model may stem from a difference in focus. The convolutional kernels prioritized understanding local information, whereas comprehending RNA sequences required an exploration of the entire sequence’s global content and meaning, as well as the overarching relationship between siRNA and mRNA.

#### Performance and Analysis on Our In-house Dataset

We evaluated the robustness and generalization of all six models on our in-house dataset. As shown in Fig 4, our siRNADesign outperformed the other models and achieves a PCC of 0.565, SPCC of 0.626 and AUC of 0.815.

**Fig. 4.**
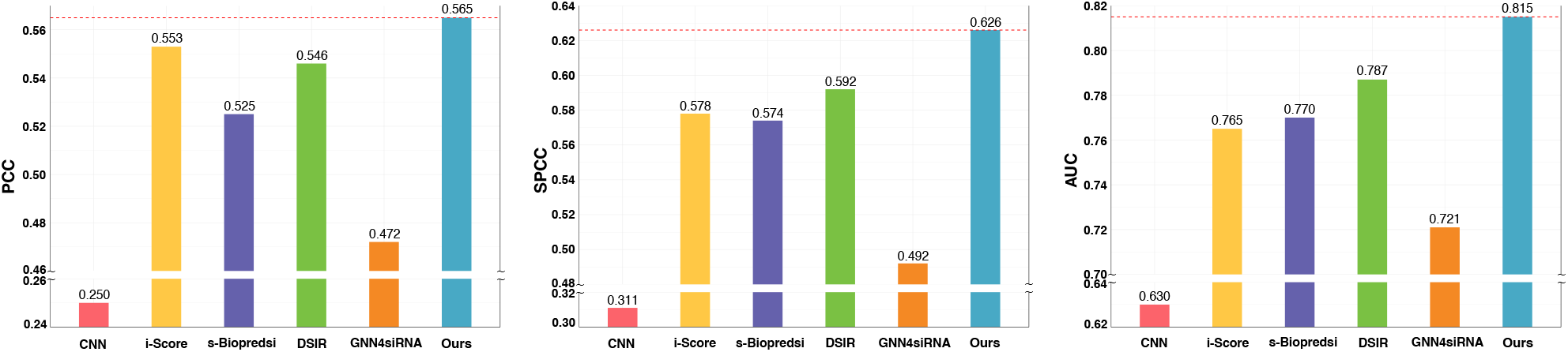
Performance of siRNADesign on our In-house Dataset. Notably, our siRNADesign significantly surpasses other methods by large performance gains, demonstrating the strong efficacy prediction capacity of our model.

Additionally, due to the limitation on the number of siRNAs, we selected top-ranked siRNAs from predictions of each method. We found that our siRNADesign had the highest proportion of siRNAs with truth efficacy above 70% (included 70%), reaching 91.7% in the top 200 and 87.5% in top 300, respectively shown in Figure 5 and Figure 6.

**Fig. 5.**
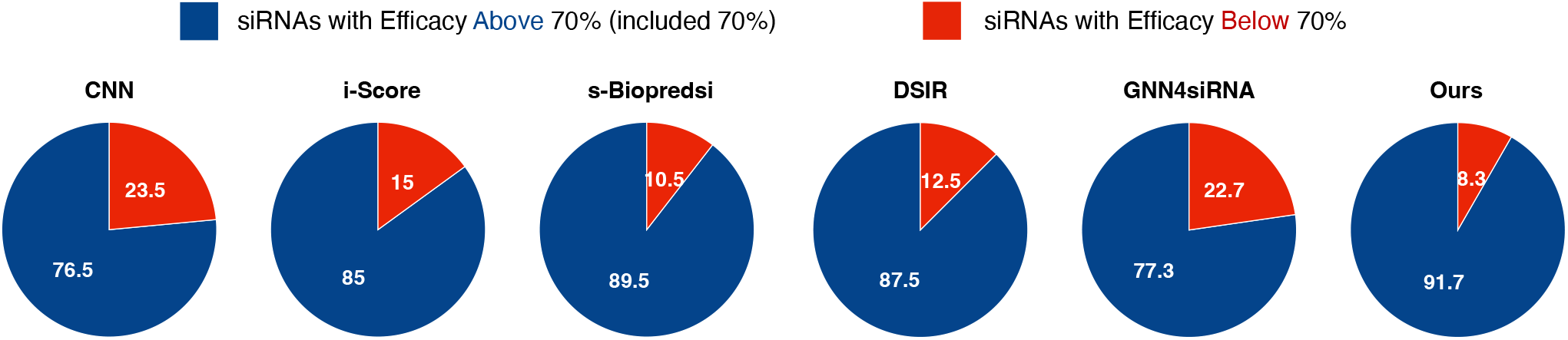
Proportion of <u>Top-200</u> siRNAs with True Efficacy Above/Below 70%. We filter the siRNAs from various models based on their top-200 predicted values **trained on siRNA-split data**. We calculate and display the percentage of these siRNAs whose actual efficacy surpasses or falls below the 70% threshold.

**Fig. 6.**
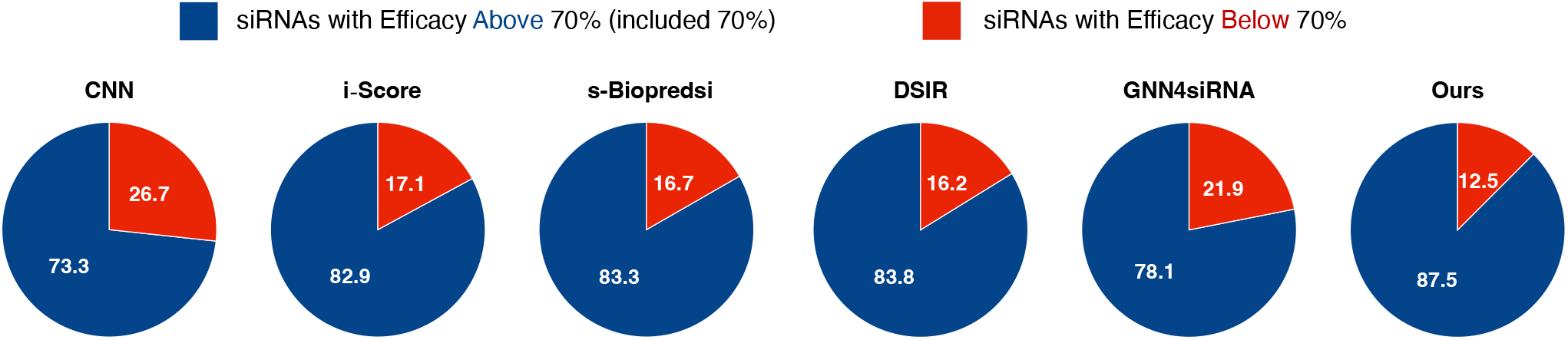
Proportion of <u>Top-300</u> siRNAs with True Efficacy Above/Below 70%. We filter the siRNAs from various models based on their top-300 predicted values **trained on siRNA-split data**. We calculate and display the percentage of these siRNAs whose actual efficacy surpasses or falls below the 70% threshold.

### Ablation study

In our ablation study, we conducted a systematic evaluation to ascertain the impact of various features on performance metrics, specifically focusing on the PCC across eight distinct categories of features on siRNA-split data. These categories encompassed sequence embeddings, thermodynamic stability, interaction probabilities, RNA-protein interactions, positional embeddings, nucleotide frequencies, rule-based codes, and G/C content percentages. As illustrated in Table 4, the integration of all features yielded the highest performance compared to other configurations. We observed a progressive enhancement in PCC as additional features were incrementally incorporated into the model. Notably, the inclusion of nucleotide frequencies resulted in the most significant improvement, with a PCC increase of 0.03.

**Table 4.** Ablation Study on Different Features Components. We report the PCC of siRNADesign on siRNA-split Data. Clearly, the combination of all the proposed extracted features performs the best.

| Sequence Embeddings | Thermodynamic Stability | Interaction Probabilities | RNA-protein Interaction | Positional Embeddings | Nucleotide Frequency | Rule Codes | G/C Percentages | PCC |
| --- | --- | --- | --- | --- | --- | --- | --- | --- |
| ✓ | ✓ | - | - | - | - | - | - | 0.731 |
| ✓ | ✓ | ✓ | - | - | - | - | - | 0.735 |
| ✓ | ✓ | ✓ | ✓ | - | - | - | - | 0.736 |
| ✓ | ✓ | ✓ | ✓ | ✓ | - | - | - | 0.738 |
| ✓ | ✓ | ✓ | ✓ | ✓ | ✓ | - | - | 0.768 |
| ✓ | ✓ | ✓ | ✓ | ✓ | ✓ | ✓ | - | 0.769 |
| ✓ | ✓ | ✓ | ✓ | ✓ | ✓ | ✓ | ✓ | **0.770** |

## Discussion

In this section, we delve into further discussions on our design choices, explorations, and directions for future research.

### Sequence Embedding by One-hot

In our model, we adopted one-hot encoding for sequence embedding rather than k-mers. First, one-hot encoding provides precise and fine-grained positional information, with each nucleotide position individually encoded. In contrast, k-mers emphasize the local pattern and context within the sequence, potentially overlooking the specific position information of individual nucleotides due to the selection of k-values. Second, one-hot encoding transforms RNA sequences into two-dimensional binary matrices, offering an approachable way to be processed by deep learning models. The features generated by k-mers potentially lead to a significant increase in the feature space.

### Incorporating RNA Base-pairing Probabilities

We first consider the impact of siRNA-mRNA co-fold binding matrices, along with the self-fold binding matrices of siRNA and mRNA, on the silencing effect of siRNAs. This consideration reflects the influence of both intrinsic and mutual interactions between siRNA and mRNA on silencing efficiency. Future work will explore novel feature engineering techniques, such as employing Convolutional Neural Networks (CNNs) for dimensionality reduction or integrating these features more seamlessly into the model.

### Incorporating RNA-AGO2 Interaction Features

We pioneer the inclusion of RNA-AGO2 interaction features to enhance siRNA efficacy prediction. The rationale behind this is that the binding between RNA and AGO2 is a critical component of the siRNA knockdown process. By integrating RNA-AGO2 interaction features, we aim for the model to grasp a more comprehensive understanding of the biological process, thereby improving prediction accuracy. Our future endeavors could explore molecular docking techniques to further consider the binding affinities between nucleic acids and proteins, which could enhance the model’s predictions of siRNA knockdown efficiency.

### The Importance of Position Embeddings

A significant oversight in many methodologies is the neglect of encoding the position of siRNAs on the mRNA within their models. However, research [38] has demonstrated that the location of siRNA target sites on the mRNA could significantly influence siRNA efficacy, with siRNAs exhibiting higher mismatch tolerance at 3’-UTR locations and greater repression in coding regions when targeted regions were structurally stable and require unraveling during translation processes [39]. Some existing methods have considered siRNAs’ positional effects but are limited to simplistic one-hot encoding strategies or sparse-semantic feature matrices with low dimensions. In contrast, our approach, siRNADesign, adopts high-dimensional transformation for positional embeddings, providing effective semantics that enhance the learning capabilities of Graph Neural Networks (GNNs).

### Incorporating Empirical Rules

Empirical rules concluded by previous studies were widely used in today’s siRNA design, such as thermodynamic stability, nucleotide frequency, siRNA rules codes and G/C percentages. This features enable our model to learn effectively at sequence level, thus enhancing the predictive power of our model. In our ablation study, we found the inclusion of siRNA nucleotide frequencies from 1-mer to 5-mers resulted in the most improvement in PCC. This supports existing research [9, 34, 35] indicating that the efficacy of gene silencing is significantly influenced by the nucleotide frequencies within the siRNA sequence. One possible reason for this influence is that nucleotide combinations contribute to RNA interference processes, such as influencing RNA secondary structure formation and complementary pairing with target mRNA, although further evidence is needed to confirm these effects. Moreover, this multi-scale feature representation, from individual nucleotides to complex multinucleotide combinations, enables the capture of dynamic changes and intricate local patterns within sequences, thereby providing a comprehensive understanding of sequence dependencies and variations.

### Performance on mRNA-split Data

We conducted a rigorous evaluation on mRNA-split data through grid searches and ablation studies, where our mode displayed comparable performance to several existing methods and surpasses that of both CNN [11] and GNN algorithms [17]. The results, **shown in Supplementary Materials**, revealed no significant performance disparity between our model and established methods such as DSIR [12] or s-Biopredsi [8], all of which outperform CNN and GNN4siRNA. Notably, our model maintained consistent stability across both siRNA-split and mRNA-split data.

### Significance of Proposed siRNA-split and mRNA-split Data Evaluation

As presented in Section 2.2, we addressed the issue of data leakage observed in previous evaluations, where siRNAs from the test set might inadvertently be included in the training phase, thus undermining the evaluation’s integrity. Moreover, we critiqued the impracticality of employing the same subset for both validation and testing as seen in existing methods.

To counter these issues, we propose and will release new data-splitting methodologies based on siRNA and mRNA, previously unconsidered by researchers. This novel approach aims to set a new benchmark for future research, encouraging more rigorous and realistic evaluations.

### Directions for Future Work

In the future, we plan to explore the chemical modification of siRNAs and the integration of transformer models and other advanced deep-learning architectures. These avenues hold the promise of further enhancing the predictive accuracy and understanding of siRNA efficacy.

## Conclusion

In this paper, we introduce **siRNADesign**, a Graph Neural Network (GNN) framework that marks a significant advancement in the prediction of siRNA efficacy. To alleviate the limitations observed in previous models [17] that lack the comprehensive understanding of siRNA silencing mechanism, our model delves deeply into both the non-empirical and empirical rules-based features of siRNA and mRNA sequences. In this way, our siRNADesign comprehensively captures the intricate features and semantics of gene silencing and achieves *state-of-the-art* results on internal datasets. Significantly, siRNADesign also demonstrates superior performance compared to current methodologies in both *in vitro* wet lab experiments and an externally collected dataset. What’s more, to address the critical issue of data leakage and the impracticality of conventional validation methods, we propose an innovative dataset-splitting methodology that enhances the evaluation’s integrity and realism. This methodology, by treating siRNA and mRNA as separate entities for dataset division, ensures a robust and unbiased assessment of model performance across various datasets. Extensive experiments, encompassing widely used and external datasets, demonstrate the superiority of our **siRNADesign**, showcasing unparalleled predictive performance and robustness across different experimental settings. We aim our model for offering a solid foundation for the development of more reliable predictive models in the field of gene silencing, and aim our proposed dataset-splitting methodology for setting a new benchmark for future research, encouraging more rigorous and realistic evaluations.

