## Supplementary file for "siRNADesign: A Graph Neural Network for siRNA Efficacy Prediction via Deep RNA Sequence Analysis"

### Overview

- Section B: Experiments on mRNA-split Data.
- Section C: Ablation Study on mRNA-split Data.
- Section D: Visualizations of Hyperparameter Tuning.

### Experiments on mRNA-split Data

In this section, we present the additional experiments of our method on mRNA-split data, besides on the siRNA-split data in the main paper.

#### Training Settings and Model Settings

Similar to the experiments on siRNA-split data in the main paper, we adopted Oputna [2] to conduct hyperparameter tuning encompassing a series of trials focusing on various parameters. We list the final settings and configurations of siRNADesign on mRNA-split data in Table S1.

| Name | Values |
|---|---|
| ***Training Hyperparameter*** | |
| Batch Size | 64 |
| Learning Rate | 1e-3 |
| Loss Function | MSE |
| Epoch Number | 38 |
| ***Model Hyperparameter*** | |
| HinSAGE Layer Size | [64, 32] |
| Hop Neighbor Sample | [4, 2] |
| Dropout Rate | 0.3 |
| Dim. of Positional Embeddings | 3 |
| Dim. of mRNA Reduced Matrices | 500 |
| Dim. of siRNA Reduced Matrices | 15 |
| Dim. of Pair Reduced Matrices | 50 |

**Table S1. Trainging Hyperparameters and Model Hyperparameters of siRNADesign on mRNA-split data.** We list the best combination of the training and model hyperparameters among all trails. 'Dim.' denotes feature dimension.

#### Performance

To further verify the effectiveness of our proposed method, we presented more results of siRNADesign compared with five previous works, i.e., GNN4siRNA [3], DSIR [4], s-Biopredsi [5], i-Score [5] and CNN model [6], on Dataset_HUVK and two external datasets.

##### Performance on Dataset_HUVK

We conducted experiments and reported the performances on mRNA-split data in Figure S1. To avoid deviation from randomness, we conducted experiments under 10 distinct splits divided under different random seeds and report the average metrics and variance values. As shown in Figure S1, our siRNADesign achieved comparable performance with small variation.

##### Performance on Simone Dataset

What's more, we evaluated the effectiveness and robustness of these models on an external dataset, Simone [1], which contained 322 siRNAs. Different from the experiments in the

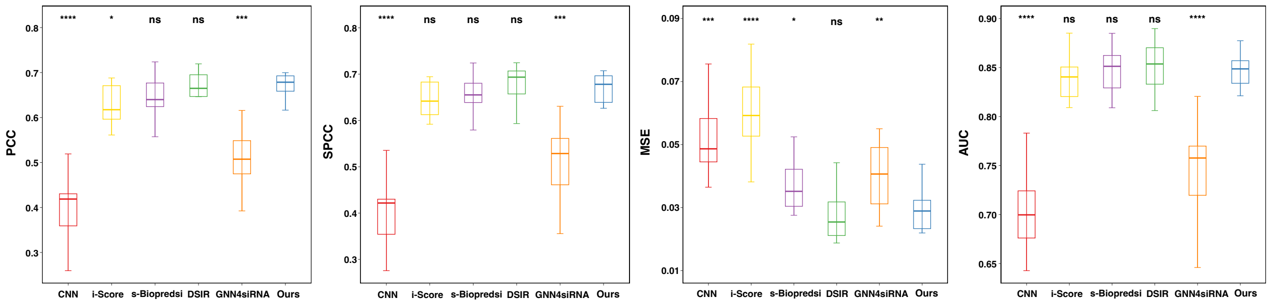


**Fig. S1. Performance of siRNADesign on mRNA-split Data (Dataset_HUVK).** Our method achieves comparable performance and remarkable stability across multiple metrics over other models. PCC: Pearson correlation coefficient; SPCC: Spearman correlation coefficient; AUC: Area Under Curve; MSE: Mean squared error. P values are calculated using paired t-test to compare the siRNADesign with the metric of other models. $* * * * P \leq 0.0001$. $* * * P \leq 0.001$. $* P \leq 0.05$. $ns\ P \geq 0.05$.

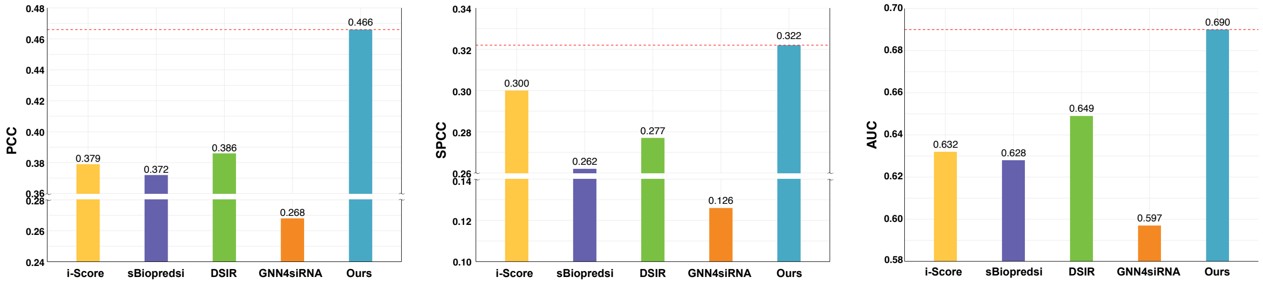


**Fig. S2. Performance of siRNADesign on an External Dataset, i.e., Simone [1].** Notably, our siRNADesign significantly surpasses other methods by large performance gains, demonstrating the strong efficacy prediction capacity of our model.

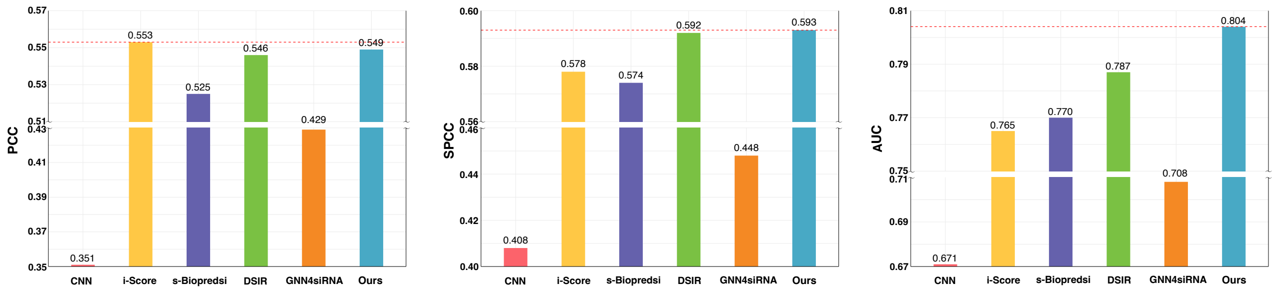


**Fig. S3. Performance of siRNADesign on our In-house Dataset.** Notably, our siRNADesign surpasses other methods by large performance gains, demonstrating the strong efficacy prediction capacity of our model.

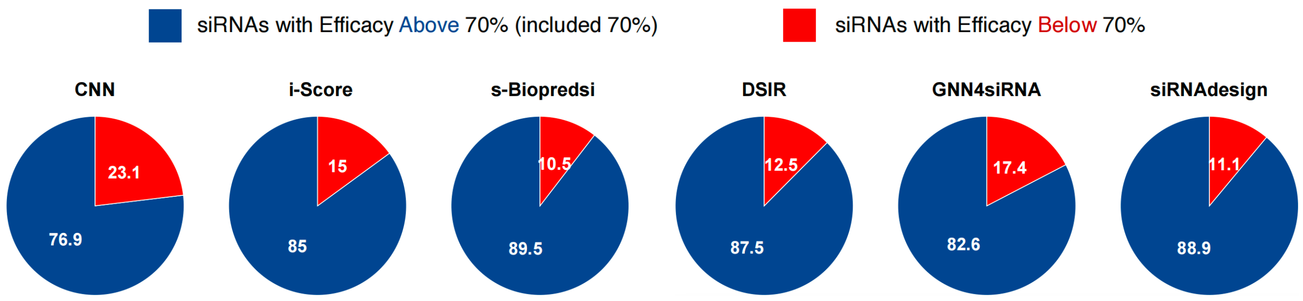


**Fig. S4. Proportion of Top-200 siRNAs with True Efficacy Above/Below 70%.** We filter the siRNAs from various models based on their top-200 predicted values **trained on mRNA-split data**. We calculate and display the percentage of these siRNAs whose actual efficacy surpasses or falls below the 70% threshold.

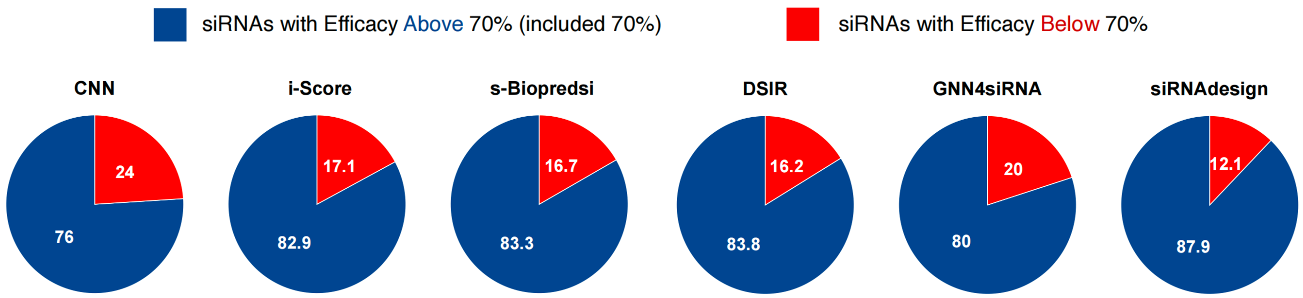


**Fig. S5. Proportion of Top-300 siRNAs with True Efficacy Above/Below 70%.** We filter the siRNAs from various models based on their top-300 predicted values **trained on mRNA-split data**. We calculate and display the percentage of these siRNAs whose actual efficacy surpasses or falls below the 70% threshold.

main paper, we first trained all models on the training set of **mRNA-split** data and utilize the pre-trained models to conduct inference on Simone. In Figure S2, we showed the evaluation results, where our siRNADesign shows the best performance across all metrics, achieving a PCC of 0.466, SPCC of 0.322 and AUC of 0.690. It further demonstrates the superior robustness of our method even for external datasets compared with other methods.

#### Performance on Our In-house Dataset

We assessed the robustness and generalization capabilities of all six models using our in-house dataset. As illustrated in Figure S3, our siRNADesign model surpassed the other models in terms of SPCC and AUC, achieving an SPCC of 0.593 and an AUC of 0.804. Additionally, our model achieved the second highest PPC at 0.549, narrowly trailing behind the leading model, i-Score [5], by only 0.004. This demonstrates the competitive performance of our siRNADesign model on mRNA-split data as well.

Additionally and similarly to the siRNA-split data evaluation, we selected top-ranked siRNAs from predictions of each method. We noticed that our siRNADesign had the competitive proportion of siRNAs with truth efficacy above 70% (included 70%), reaching 88.9% in the top 200 and 87.9% in top 300, respectively shown in Figure S4 and Figure S5.

### Ablation Study on mRNA-split Data

In this section, we extended our systematic evaluation to examine the influence of various feature categories on

| Sequence Embeddings | Thermodynamic Stability | Interaction Probabilities | RNA-protein Interaction | Positional Embeddings | Nucleotide Frequency | Rule Codes | G/C Percentages | PCC |
|---|---|---|---|---|---|---|---|---|
| ✓ | ✓ | - | - | - | - | - | - | 0.646 |
| ✓ | ✓ | ✓ | - | - | - | - | - | 0.652 |
| ✓ | ✓ | ✓ | ✓ | - | - | - | - | 0.652 |
| ✓ | ✓ | ✓ | ✓ | ✓ | - | - | - | 0.654 |
| ✓ | ✓ | ✓ | ✓ | ✓ | ✓ | - | - | 0.662 |
| ✓ | ✓ | ✓ | ✓ | ✓ | ✓ | ✓ | - | 0.669 |
| ✓ | ✓ | ✓ | ✓ | ✓ | ✓ | ✓ | ✓ | **0.672** |

**Table S2. Ablation Study on Different Features Components on mRNA-split Data.** We report the PCC of siRNADesign on siRNA-split Data. Clearly, the combination of all the proposed extracted features performs the best.

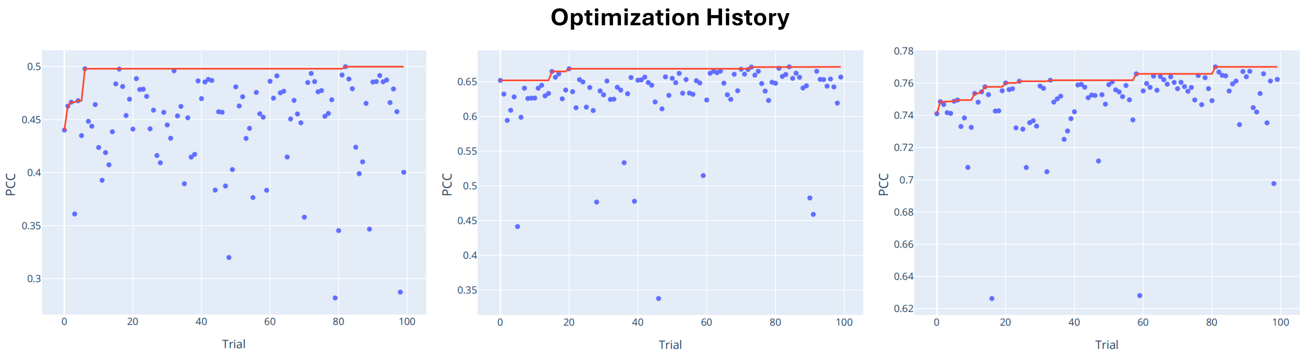


(a) **GNN4siRNA** (b) **siRNADesign** on mRNA-split Data (c) **siRNADesign** on siRNA-split Data

**Fig. S6. Optimization history of siRNADesign and GNN4siRNA models using Optuna.** In subfigures (b) and (c), the concentration of data points indicates higher stability and robustness of the siRNADesign model, with a higher density of optimal points suggesting superior optimization outcomes. This visualization effectively demonstrates the enhanced performance and reliability of our approach in computational siRNA design.

performance metrics, focusing on the PCC using mRNA-split data. The evaluated feature categories included sequence embeddings, thermodynamic stability, interaction probabilities, RNA-protein interactions, positional embeddings, nucleotide frequencies, rule-based codes, and G/C content percentages. As depicted in Figure S2, the cumulative integration of these features led to the highest performance relative to other configurations. A gradual improvement in PCC was observed as features were progressively added to the model.

### Visualizations of Hyperparameter Tuning.

In this section, we visualize the optimization history and parallel coordinate plot that illustrates the relationships and patterns across multiple variables, allowing for an in-depth analysis of trends and anomalies in the datasets.

#### Optimation History

In Figure S6, we presented the optimization history of our siRNADesign and GNN4siRNA [3] model (a) using Optuna [2], a sophisticated hyperparameter optimization framework. Subfigures (b) and (c) depicted the optimization trajectories on both mRNA-split and siRNA-split datasets, highlighting the performance and robustness of our models. Notably, subfigures (b) and (c), which represent our siRNADesign model, showed a more concentrated distribution of data points. This clustering indicated that our model not only achieved more consistent performance across different datasets but also exhibited superior robustness. Furthermore, the density of optimal points in these subfigures was significantly higher, suggesting that our model was capable of reliably reaching peak performance levels, underscoring its effectiveness in siRNA design. This visual analysis served as a clear demonstration of the practical advantages our model holds over existing approaches, as facilitated by the Optuna optimization engine.

#### Parallel Coordinate Plot

In Figure S7, we used parallel coordinate plots to visualize the optimization of our siRNADesign model on both mRNA-split and siRNA-split data, shown in subfigures (a) and (b). This visualization included a range of trials examining hyperparameters such as the sizes of HinSAGE layers, hop neighbor samples, batch sizes, dropout rates, dimensions of positional embeddings, and dimensions of reduced matrices for both RNA base-pairing probabilities and siRNA-mRNA base-pairing probabilities. The plot indicated that our model's performance was highly sensitive to changes in batch size and epochs, suggesting that these parameters were pivotal for achieving optimal results. Conversely, the model demonstrated a lower sensitivity to other hyperparameters like dropout rates and dimensions of positional embeddings. This lower sensitivity could signify a robustness in the model's architecture, implying that it could maintain effective performance under varying conditions of less critical parameters. Such robustness was advantageous in practical applications where varying data conditions and computational constraints required a model to perform consistently without extensive re-tuning of all parameters.

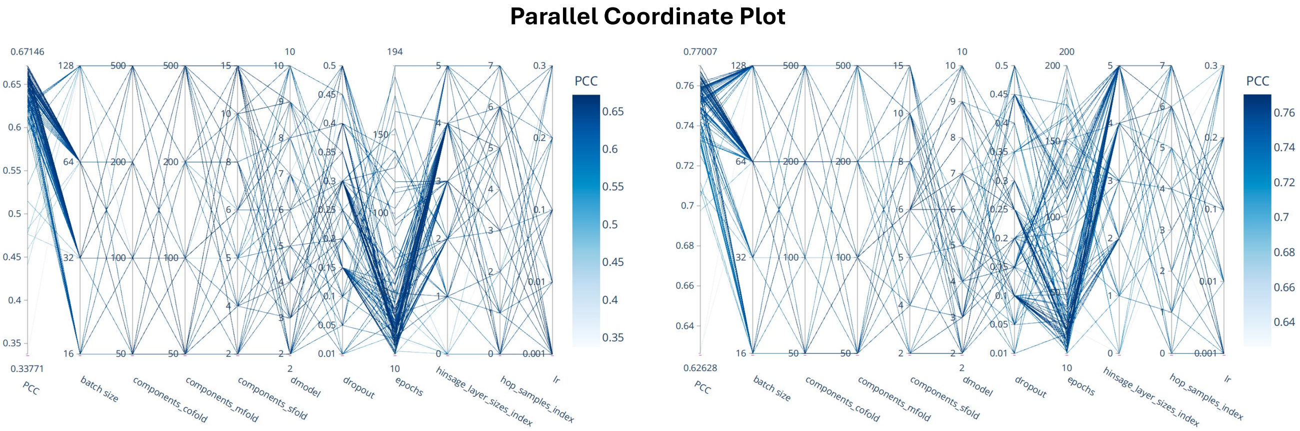


(a) siRNADesign on mRNA-split Data

(b) siRNADesign on siRNA-split Data

**Fig. S7. Hyperparameter sensitivity analysis of the siRNADesign model visualized through a Parallel Coordinate Plot.** Subfigures (a) and (b) illustrate the model's response to varying parameters on mRNA-split and siRNA-split data. The plot highlights the model's significant sensitivity to batch size and epochs, while showing robustness to variations in other parameters like dropout rates and embedding dimensions.
